# Perturbations of glutathione and sphingosine metabolites in Port Wine Birthmark patient-derived induced pluripotent stem cells

**DOI:** 10.1101/2023.07.18.549581

**Authors:** Vi Nguyen, Jacob Kravitz, Chao Gao, Marcelo L. Hochman, Dehao Meng, Dongbao Chen, Yunguan Wang, Anil G. Jegga, J Stuart Nelson, Wenbin Tan

## Abstract

Port Wine Birthmark (PWB) is a congenital vascular malformation in the skin, occurring in 1-3 per 1,000 live births. We recently generated PWB-derived induced pluripotent stem cells (iPSCs) as clinically relevant disease models. The metabolites associated with the pathological phenotypes of PWB-derived iPSCs are unknown, which we aimed to explore in this study. Metabolites were separated by ultra-performance liquid chromatography and were screened with electrospray ionization mass spectrometry. Orthogonal partial least-squares discriminant analysis, multivariate and univariate analysis were used to identify differential metabolites (DMs). KEGG analysis was used for the enrichment of metabolic pathways. A total of 339 metabolites were identified. There were 22 DMs confirmed with 9 downregulated DMs including sphingosine and 13 upregulated DMs including glutathione in PWB iPSCs as compared to controls. Pathway enrichment analysis confirmed the upregulation of glutathione and downregulation of sphingolipid metabolism in PWB-derived iPSCs as compared to normal ones. We next examined the expression patterns of the key factors associated with glutathione metabolism in PWB lesions. We found that hypoxia-inducible factor 1α (HIF1α), glutathione S-transferase Pi 1 (GSTP1), γ-glutamyl transferase 7 (GGT7), and glutamate cysteine ligase modulatory subunit (GCLM) were upregulated in PWB vasculatures as compared to blood vessels in normal skins. Our data demonstrate that there are perturbations in sphingolipid and cellular redox homeostasis in the PWB vasculature, which may facilitate cell survival and pathological progression. Our data imply that upregulation of glutathione may contribute to laser-resistant phenotypes in the PWB vasculature.

## Background

Congenital capillary malformation, also known as port-wine birthmarks or stains (PWB or PWS), is one of the most common types of congenital vascular malformations (CVMs). PWB result from developmental defects in the skin vasculature with an estimated prevalence of 1-3 per 1,000 live births [1]. This skin lesion can be isolated or in association with other CVMs in children, including arteriovenous malformations, Parkes-Weber syndrome, and Sturge-Weber syndrome (SWS) [2]. For example, SWS, an ipsilateral leptomeningeal angiomatosis syndrome, has been reported in approximately 25% of infants with forehead PWB lesions [3]. At birth, PWBs appear as flat red macules; lesions progressively darken to purple during the growth of afflicted children. By middle age, lesions in many patients show a development of vascular nodules that are susceptible to spontaneous bleeding or hemorrhage [1]. Moreover, a PWB child’s quality of life is greatly affected during their development and growth due to devastating lifelong psychological and social impacts [1].

The vascular phenotypes of PWB lesions typically exhibit progressive dilatation of dermal capillaries invloving the entire skin, which are characterized by proliferation of endothelial cells (ECs) and smooth muscle cells (SMCs), replication of basement membranes, and disruption of vascular barriers [4, 5]. Pathologically, PWB ECs are immature and differentiation-defective, expressing biomarkers of stem cells and endothelial progenitor cells (EPCs) [4, 6]. The pulsed dye laser (PDL) is the treatment of choice for PWB. Unfortunately, less than 10% of patients achieve complete removal of PWB after multiple PDL treatments [7]. Moreover, approximately 20% of lesions show little or no response to laser exposures, i.e., laser-resistant [8]. Re-darkening of PWB lesions can occur in 16.3%-50% of patients within five years after multiple treatments [8]. A recent review showed that there has been little improvement in clinical outcomes using laser-based modalities for PWB treatment over the past three decades [9]. We posit that the differentiation-defective PWB ECs are responsible for these clinical manifestations. For example, PWB ECs likely survive and proliferate after laser treatments, resulting in revascularization of lesional blood vessels [4, 10].

The absence of clinically relevant cell and animal models has been a long-term obstacle to understand the pathogenesis and to develop therapeutics for PWB. In an effort to overcome this barrier, we recently generated PWB patient-derived induced pluripotent stem cells (iPSCs) by introducing the “ Yamanaka factors” (Oct3/4, Sox2, Klf4, c-Myc) [11] into lesional dermal fibroblasts. The iPSCs were differentiated into clinically relevant lesional ECs [12]. These iPSC-derived ECs recapulated many pathological phentoypes of PWB vasculatures, including formation of enlarged vasculatures *in vitro* and *in vivo* and impairments of Hippo and Wnt pathways [12]. However, the metabolites reflecting the pathological phenotypes of PWB-derived iPSCs are unknown. Identification of metabolic signatures from PWB-derived iPSCs will provide us with new insights into causing signalosomes that lead to the developmental defects in lesional vasculature.

## Methods

### Tissue preparation

The study (#1853132) was approved by the Institutional Review Board at the Prisma Health Midlands. De-identified surgically excised hypertrophic and nodular PWB lesions (n=4) and de-identified surgically discarded normal skin tissues (n=5) were collected during this study or through our previous studies [4, 5, 13].

### PWB iPSC culture and sample preparation

Generation of the PWB iPSC and control iPSC lines from normal skins was detailed in our recent study [12]; these cell lines were maintained and propagated under feeder-free conditions using Essential 8 Medium on a geltrex-coated plate (ThermoFisher, Waltham, MA). The following iPSC lines were used in this study: PWB_4221_3, PWB_4221_6, PWB_3921_9, PWB_3921_16, Control_52521_8, and control_52521_9 [12]. There were experimental duplications for PWB_4221_3, PWB_3921_9, and Control_52521_8, resulting in a total of 9 samples. The iPSCs (∼10^6^ cells) were dissociated using StemPro Accutase (ThermoFisher, Waltham, MA); cell pellets were collected and stored at -80 °C until processing. Samples were thawed with the addition of 300 ul of 80% methanol. Samples were vortexed for 60 seconds, sonicated for 30 minutes at 4°C, and incubated at -20°C for one hour. The samples were then subjected to centrifugation at 12,000 rpm at 4°C for 15 minutes. The supernatant (200 µl) was mixed with 5 µl of L-o-Chlorophenylalanine (0.14 mg/ml) for liquid chromatography mass spectrometry (LC-MS) analysis.

### Metabolomics by LC-MS

For metabolomics analysis by LC-MS, separation was carried out using the Vanquish Flex Ultra-performance liquid chromatography (UPLC) along with the Q Exactive MS (ThermoFisher, Waltham, MA), and it was screened with electrospray ionization MS (ESI-MS). The LC system comprised the ACQUITY UPLC HSS T3 (100×2.1 mm×1.8 μm) with UPLC. The mobile phase was composed of 0.05% formic acid water and acetonitrile with gradient elution (0-1 min, 5% acetonitrile; 1-12 min, 5%-95% acetonitrile; 12-13.5 min, 95% acetonitrile; 13.5-13.6 min, 95%-5% acetonitrile; 13.6-16 min, 5% acetonitrile). The flow rate of the mobile phase was 0.3 mL/min. The column temperature was maintained at 40°C, and the sample manager temperature was set at 4°C. MS in ESI+ and ESI-mode were listed as follows: (1) ESI+: Heater Temp 300°C; Sheath Gas Flow rate, 45 arb; Aux Gas Flow Rate, 15 arb; Sweep Gas Flow Rate, 1 arb; spray voltage, 3.0 KV; Capillary Temp, 350°C; S-Lens RF Level, 30%; and (2) ESI-: Heater Temp 300°C, Sheath Gas Flow rate, 45 arb; Aux Gas Flow Rate, 15 arb; Sweep Gas Flow Rate, 1 arb; spray voltage, 3.2 KV; Capillary Temp,350°C; S-Lens RF Level, 60%.

### Immunohistochemistry (IHC)

Skin biopsies were fixed in 4% buffered paraformaldehyde and embedded in paraffin. The paraffin sections (6 µm thickness) were deparaffinized and antigen retrieval was performed in 10 mM sodium citrate buffer (pH 6.0) at 97^°^C for 2 hrs. Sections were then incubated in a humidified chamber overnight at 4^°^C with the following primary antibodies: anti-hypoxia-inducible factor 1α (HIF1α) (Santa Cruz Biotech., Dallas, TX, USA; #SC-10790; 1:100 dilution); anti-glutathione S-transferase Pi 1 (GSTP1) (Proteintech, #15902-1-AP; 1:1000 dilution), anti-γ-glutamyl transferase 7 (GGT7) (Proteintech, Rosemont, IL, USA; #24674-1-AP; 1:1000 dilution), and anti-glutamate cysteine ligase modulatory subunit (GCLM) (Proteintech, #14241-1-AP; 1:1000 dilution). Biotinylated anti-mouse or -rabbit secondary antibodies were incubated with sections for 2 hrs at room temperature after the primary antibody reaction. An indirect biotin avidin diaminobenzidine (DAB) system (Dako, Glostrup, Denmark) was used for detection. IHC scores were developed as previous reported [13] and tailored for individual blood vessels using the following two factors: (1) an intensity factor ranging from 0-5, which was the average intensity of all ECs in one blood vessel; Each EC was graded as the following categories: no staining, 0; very weak staining, 1; mild staining, 2; intermediate staining, 3; strong staining, 4; and very strong staining, 5; (2) a percentage factor ranging from 0-5 as well, which is equal to multiplying the percentage (%) of immunoreactive positive ECs in one blood vessel by a factor of 5. For each blood vessel, an antibody immunoreactivity score was estimated by multiplying the intensity and percentage factors, ranging from 0 to 30.

### Statistical analysis

The raw data were acquired and aligned using the Compound Discoverer (3.0, Thermo) based on the m/z value and the retention time of the ion signals. Ions from both ESI- and ESI+ were merged and imported into the SIMCA-P program (version 14.1) for multivariate analysis. Principal Components Analysis (PCA) was used for data visualization and outlier identification. Supervised regression modeling was then performed on the data set using Partial Least Squares Discriminant Analysis (PLS-DA) or Orthogonal Partial Least Squares Discriminant Analysis (OPLS-DA) to identify the potential biomarkers. The biomarkers were filtered and confirmed by combining the results of the Variable Importance in Projection (VIP) values (VIP > 1.5) and t-test values (p < 0.05). The quality of the fitting model was explained by R2 and Q2 values. R2 displayed the variance explained by the model and indicated the quality of the fit. Q2 displayed the variance in the data, indicating the model predictability.

### Results

The total ions chromatograms (TIC) showed the summing up intensities of all mass spectral peaks associated metabolites among samples (Figs 1A and B; Supplementary Figs 1 and 2). The data were normalized after alignment. PCA showed that the quality control samples were highly clustered, but no clear grouping trend was observed between the PWB and control groups (Supplementary Figs 3A and 3B). To eliminate any non-specific effects and confirm the biomarkers, PLS-DA or OPLS-DA was used to compare metabolic changes in the two groups, respectively. Both analyses showed a clear separation of the PWB and control groups (Figs 1C and D; Supplementary Figs 4 and 5). There were 339 detected metabolites amongst the samples, with 156 metabolites detected in the ESI-mode, 201 in the ESI+ mode, and 18 overlapping metabolites detected in both modes.

**Fig 1.**
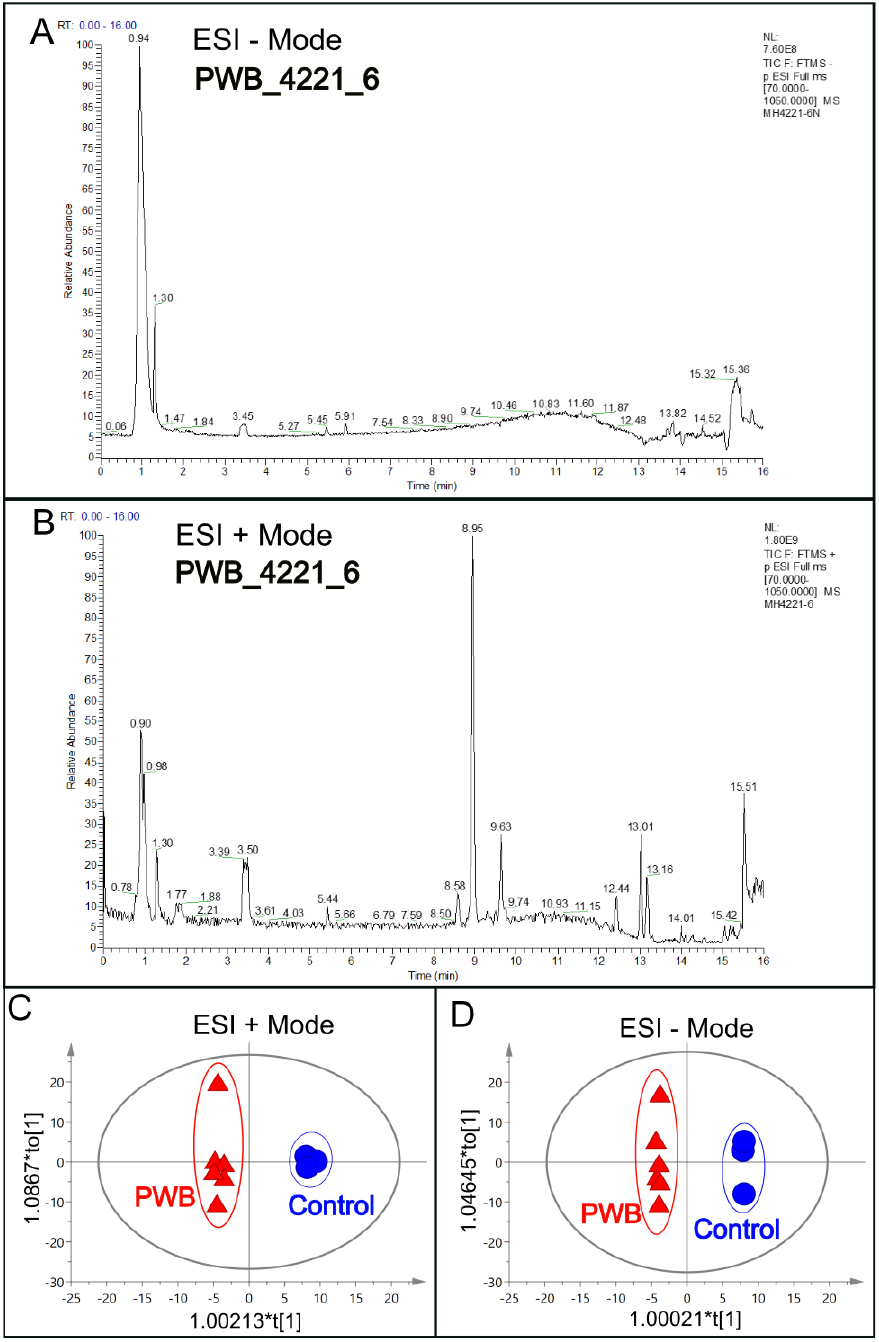
An example of total ions chromatogram (TIC) representing the summed intensity with all detected mass spectral peaks associated with metabolites. A, TIC from the PWB_4221_6 iPSC line in ESI-mode; B, TIC from the same iPSC line in ESI+ mode; C, OPLS-DA model showing scattering scores and cluster tendencies among all samples in ESI + mode; D, OPLS-DA model showing scattering scores and cluster tendencies among all samples in ESI - mode. Digits on the X or Y axis are eigenvalues of regression coefficient for the predictive principal component (X) or the orthogonal component (Y), respectively.

Significantly changed metabolites between the two groups were called out based on VIP scores (VIP > 1.5) (Supplementary Figs 6). The metabolic biomarkers were filtered and collected by combining the results of VIP scores (VIP > 1.5) and t-test values (p < 0.05) (Figs 2A-B). The significantly differential metabolites (DMs) (absolute log_2_(fold change) > 0.5, and p value < 0.05) were determined using univariate analysis (Figs 2C-D). The final metabolic biomarkers were confirmed using the data of accurate masses and MS/MS fragments with the criteria of VIP score > 1.5, absolute log_2_(fold change) > 0.5, and p value < 0.05. In particular, there was an upregulation of glutathione and downregulation of sphingosine in PWB iPSCs as compared to control ones. The representative mass spectra for glutathione and sphingosine were shown in Fig 3 and supplementary table 1. A total of 22 metabolic biomarkers were identified in the final list (8 in ESI - and 16 in ESI + with 2 overlapped ones) (table 1, Figs 4A-B). Next, the metabolic network was revealed using KEGG enrichment analysis (Figs 4C-D). Significantly affected metabolic pathways included sphingolipid, glutathione, arachidonic acid, pyrimidine, pentose and glucuronate interconversions, galactose, ascorbate and aldarate, starch and sucrose, amino sugar and nucleotide sugar, alanine, aspartate, and glutamate.

**Fig 2.**
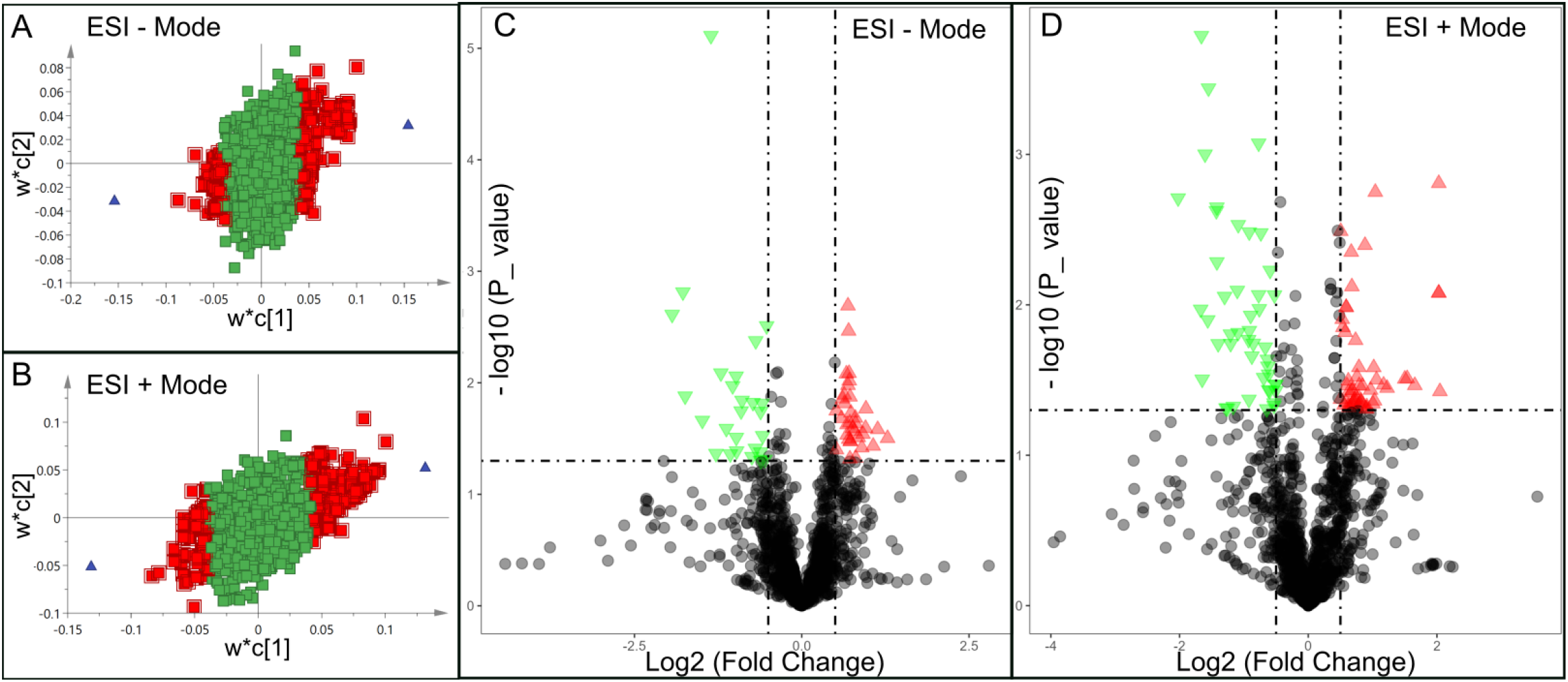
Discovery of differential metabolites (DMs) by multivariate analysis. A and B, distribution of significant metabolites detected in ESI – (A) and ESI + (B) modes using a PLS-DA model resulting in coefficients for the variables in a w*c loading plot. w, PLS-weights for the X-variables; c, PLS-weights for the Y-variables; Red box, metabolites with VIP > 1.5; green box: metabolites with VIP < 1.5; C and D, volcano plots showing clusters of DMs detected in ESI – (C) and ESI + (D) modes. Blue triangle, significantly downregulated metabolites; red triangle: significantly upregulated metabolites; black circle: insignificant metabolites; vertical dash lines: log_2_ (fold change) = ± 0.5; horizontal dash line: p value = 0.05.

**Fig 3.**
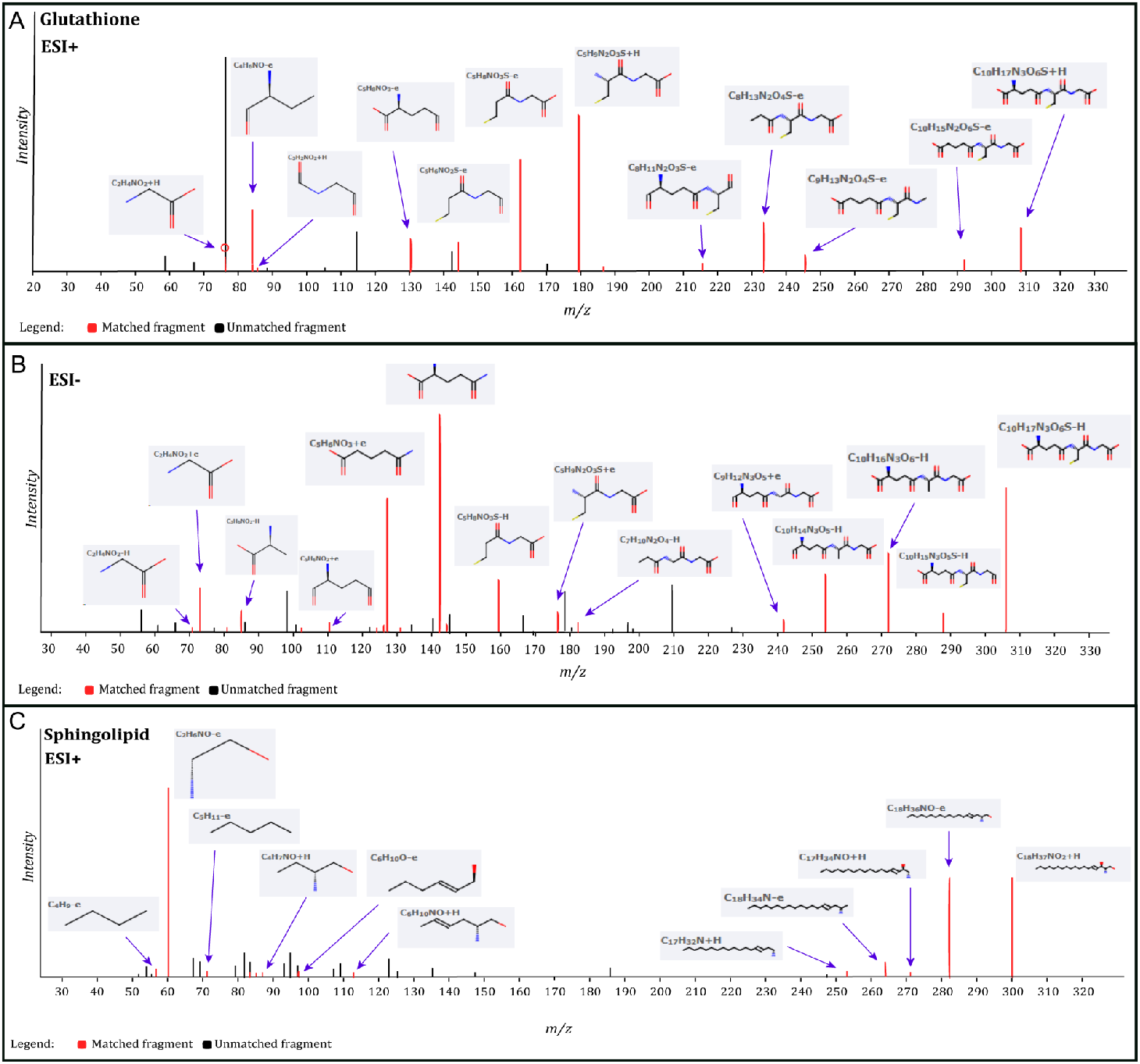
Identification of glutathione and sphingosine metabolites. A, identified matched glutathione MS fragments in ESI + mode; B, identified matched glutathione MS fragments in ESI - mode; C, identified matched sphingosine MS fragments in ESI + mode. The MS chromatograms of glutathione and sphingosine fragments from each sample were extracted and aligned into one chromatogram.

**Fig 4.**
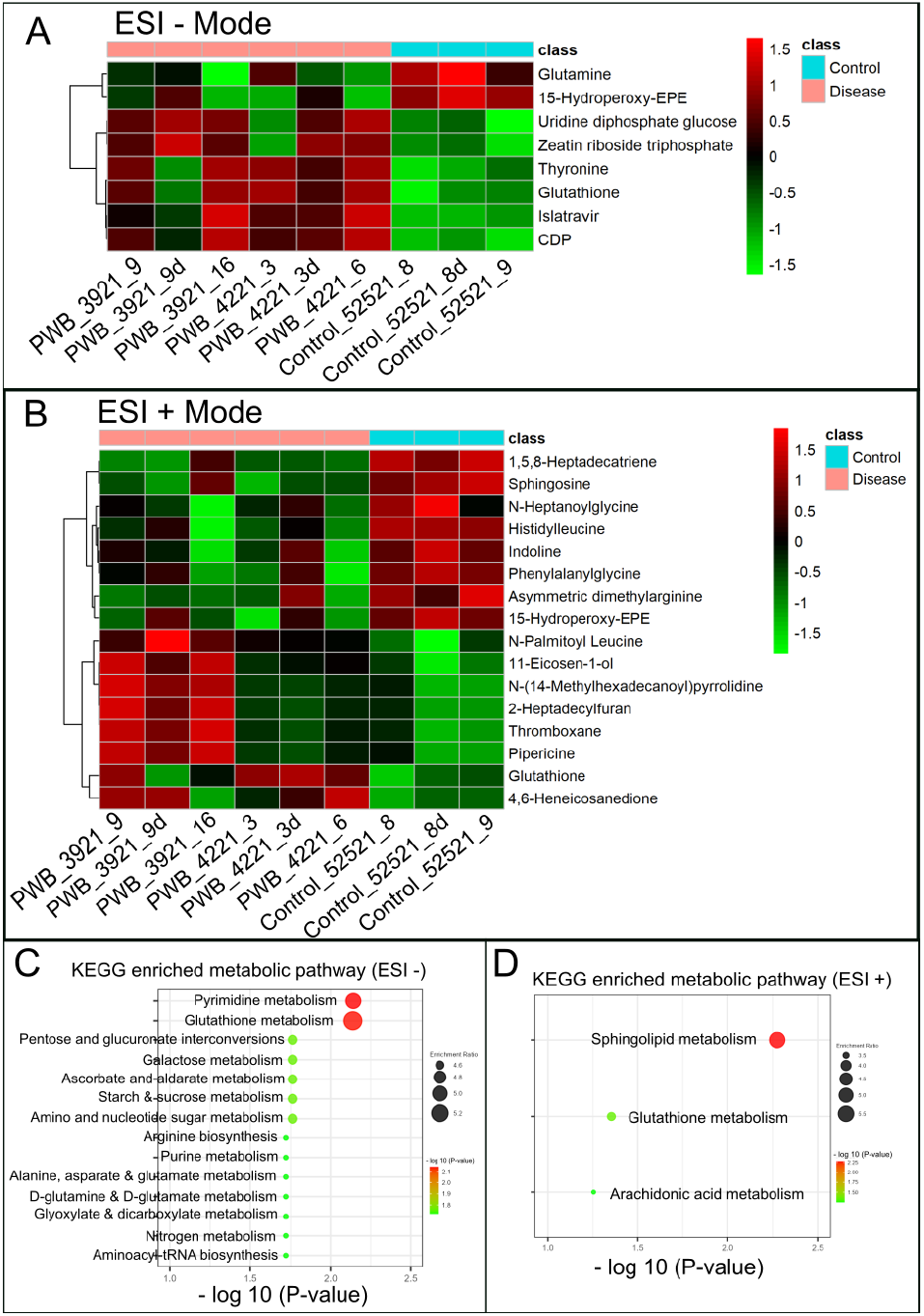
Hierarchical cluster analysis (HCA) and KEGG pathway enrichment of metabolome data. A & B, the heatmap of significantly DMs identified in ESI – (A) and ESI + mode (B) in PWB iPSCs as compared to the control ones; HCA was performed using the complete linkage algorithm of the program Cluster 3.0 and the results are visualized using Pheatmap 1.0.12 (Raivo Kolde). C & D, enriched KEGG pathways related to the perturbed metabolic networks involving DMs identified in ESI – (C) and ESI + mode (D); the enriched metabolic pathways are indicated by color (-log_10_ (P value)) and size (black ball, enrichment ratio). PWB_3921_9d, PWB_4221_3d, and Control_52521_8d were experimental duplications of PWB_3921_9, PWB_4221_3, and Control_52521_8, respectively.

**Table 1.**
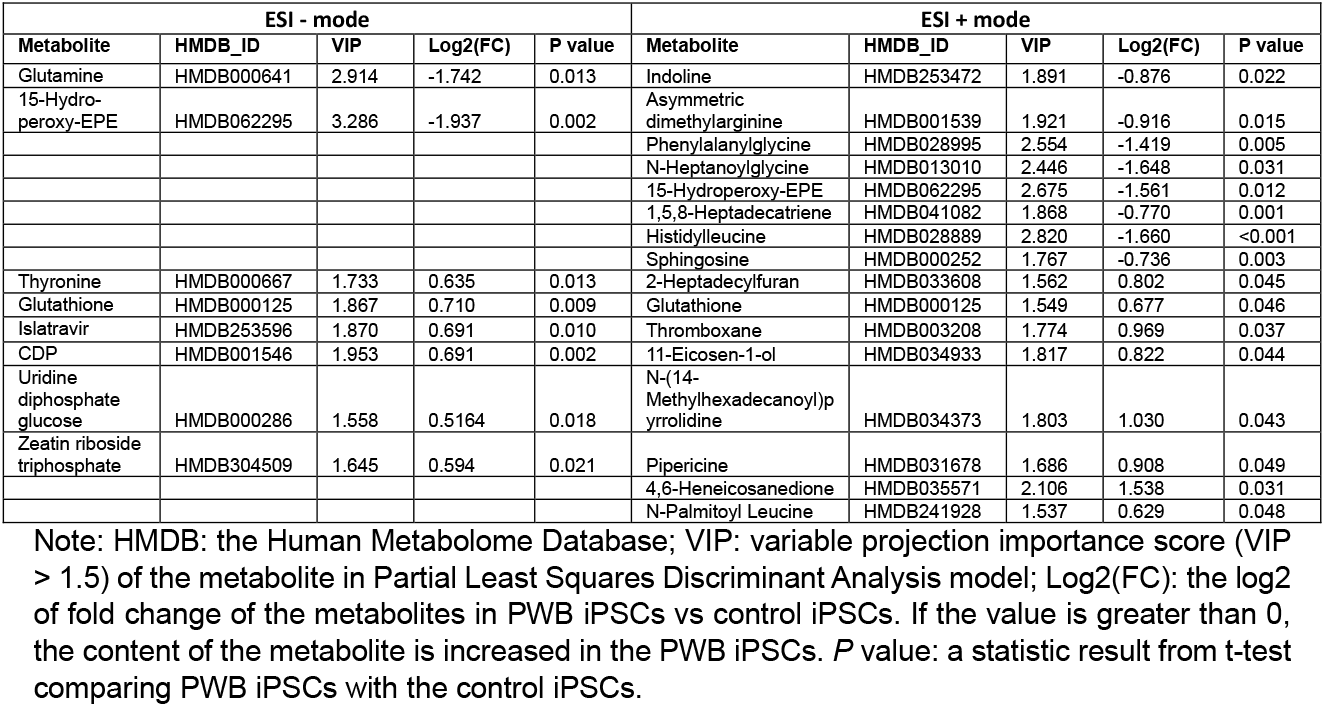
DMs in PWB iPSCs as compared to control iPSCs.

We next focused on the glutathione pathway for further validation in PWB lesions because a high level of glutathione could directly scavenge laser-induced free radicals, such as reactive oxygen species (ROS) and nitric oxide (NO) [14, 15], thus contributing to the laser resistance observed in some patients. We posited that a status of persistent mild hypoxia in PWB lesions may act as the driving force to increase glutathione, which could be indicated by an elevated level of HIF1α. Therefore, we performed an IHC to examine the expression patterns of HIF1α and several key enzymes related to glutathione metabolism in PWB lesions, including GSTP1, GGT7, and GCLM. The ECs in normal skins showed mild or moderate staining signals for the antibodies examined (Fig 5), which were also consistent with the IHC data on human skins from the human protein atlas (https://www.proteinatlas.org/). In PWB lesions, these antibodies showed moderate to strong immunoreactive signals on ECs (Fig 5, Supplementary Figs 7). The immunoreactive scores of these antibodies were significantly higher in PWB vasculatures than in normal dermal blood vessels (Fig 5).

**Fig 5.**
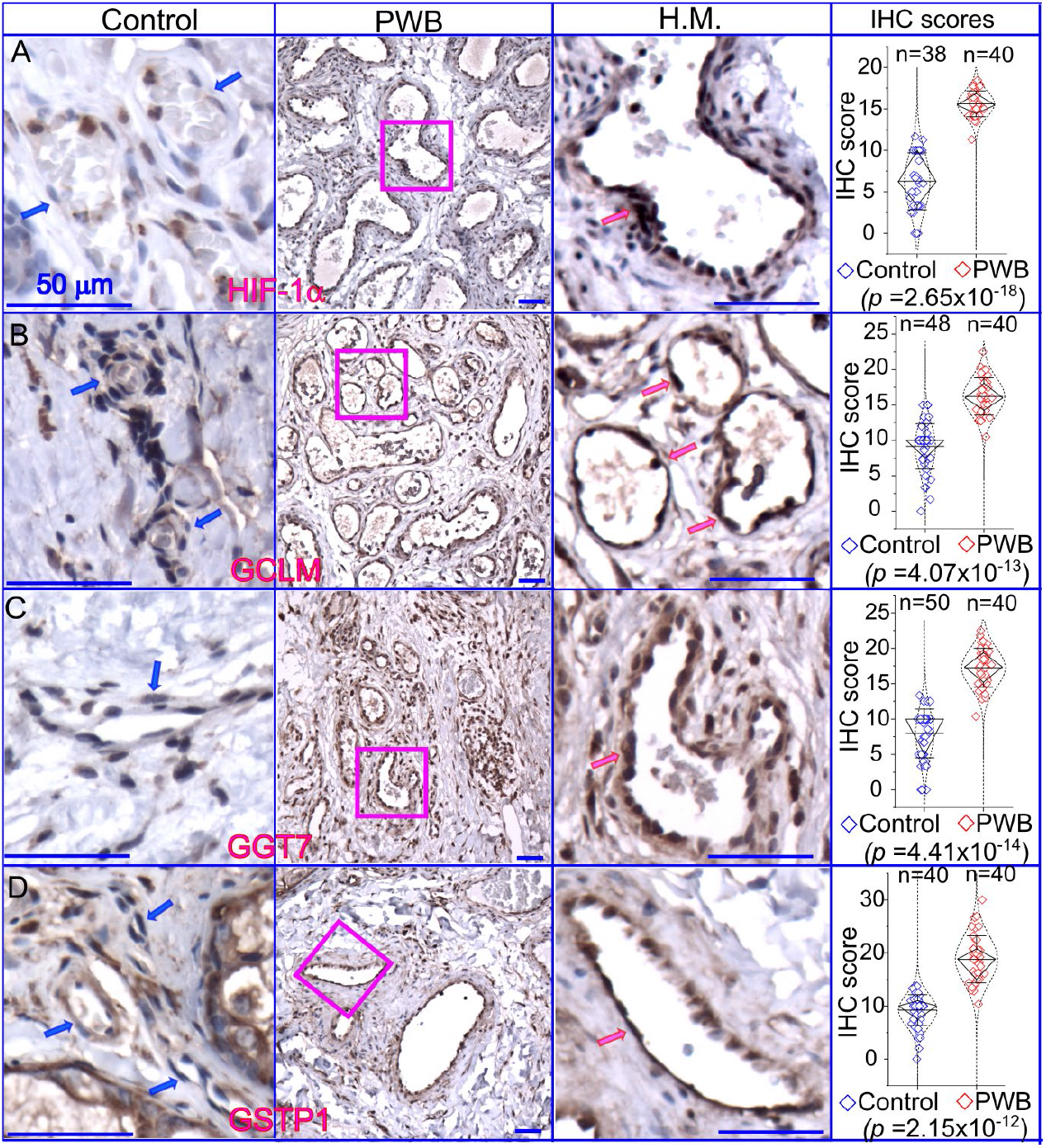
Expressions of key factors associated with glutathione metabolism in PWB lesions. The IHC assays by using antibodies recognizing HIF-1α (A), GCLM (B), GGT7 (C), and GSTP1 (D) show the immunoreactive blood vessels in control skin or PWB lesions. H.M., a higher magnification from the pink boxed area in the left panel showing immunoreactive positive PWB blood vessels for the corresponding antibodies. Scale bar: 50 µm. n, number of blood vessels from PWB (4 subjects) or normal ones (5 subjects). Whiskers: mean±S.D.; Diamond boxes: IQR; Dotted curves: data distribution. Paired *t*-test was used for comparing arbitary IHC scores. Blue arrows: control dermal capillaries; red arrows: PWB blood vessels.

## Discussion

In this study, we have identified that sphingosine is downregulated, and glutathione is upregulated, in PWB iPSCs as compared to normal controls through a non-targeted metabolomic study, which is the first of its kind. PWB vasculatures have elevated HIF-1α levels, demonstrating persistent and mild hypoxia as the driving force to increase glutathione. The dysregulation of crucial enzymes involved in glutathione metabolism in PWB lesions further supports our metabolic data. Our study provides the first link between metabolic biomarkers such as sphingosine and glutathione and the pathological progression of PWB, opening new avenues of investigations in the field.

Ceramide, sphingosine, and their derived sphingolipids are active lipids that execute diverse cellular signaling for dynamic functions [16, 17]. Ceramide is produced by breakdown of metabolically inactive membrane sphingomyelin by sphingomyelinase or *de novo* synthesis. In contrast, sphingosine is generated through de-acylation of ceramide by ceramidase. Sphingosine can further generate sphingosine-1-phosphate (S1P) by sphingosine kinases. Sphingosine has been shown to have multiple biological functions, including pro-apoptosis, cell cycle arrest, cytoskeleton regulation, endocytosis, autophagic process, and antimicrobial effects [18-20]. Sphingosine is considered as an endogenous inhibitor of protein kinase C [20-22] and also involves the induction of apoptosis by activation of PKA and inhibition of MAPK [16, 17]. Our previous studies have shown activation of PKC, MAPK, and PI3K pathways in the PWB vasculature [13, 23]. These data suggest that there are two possible mechanisms by which sphingosine contributes to the pathological development of PWB. First, downregulated sphingosine in PWB lesions may cause less inhibition of PKC and MAPK, leading to cell proliferation. Second, a lower level of sphingosine may attenuate PDL-induced apoptosis in the lesional vasculature due to its pro-apoptotic function. However, the detailed mechanisms are yet to be determined. Nevertheless, our data first provide a potential link between sphingolipids and the pathogenesis of PWB.

Our data demonstrate that glutathione is elevated in PWB iPSCs. Glutathione is the most abundant nonprotein thiol in the cell and plays diverse roles in antioxidant defense, nutrient metabolism, and cellular signaling [24-26]. Glutathione plays major beneficial roles in maintaining cellular redox homeostasis by acting as a free radical scavenger and a detoxifying agent; it participates in and contributes to multiple cellular processes including proliferation, cell division, and differentiation [25, 26]. The synthesis of glutathione is affected and regulated by many metabolic pathways, including glutamine, cysteine, glycine, glutamate, pentose, etc.[27-29] Our KEGG data showed that many of these pathways are dysregulated in PWB iPSCs. In addition, growth factors, commons stressors, and cellular metabolic reprogramming also regulate intracellular glutathione levels [25, 27, 30, 31]. Alternatively, dysregulated glutathione metabolism has been shown to have a pathogenic role in malignancy [25]. Emerging evidence has shown that glutathione levels are increased in tumor cells to help them survive in response to an accumulation of large amounts of ROS production [32-34]. Excessive glutathione promotes tumor progression, correlates with increased metastasis, and increases chemo-therapeutic resistance [25, 35-39]. We hypothesize that elevated glutathione plays both beneficial and pathogenic roles in PWB lesions. The progressively dilated PWB vasculature is a low-flow vascular malformation, which results in persistently mild hypoxia in lesions and is supported by elevated levels of HIF-1α in PWB vasculature. It is reasonable to posit that there are moderately increased ROS levels in PWB lesions under such mild hypoxic conditions. Moderate ROS levels will result in vascular cell survival and proliferation by activating signaling pathways including MAPK, JNK, PKC, and PI3K/AKT. Therefore, increased glutathione, through upregulation of its metabolic enzymes such as GGLM, GGT7, and GSTP1, will be needed for PWB vascular cells to scavenge excessive oxidation and maintain an intricate antioxidant status to survive. Also, elevated glutathione levels will help lesional vasculature survive after laser treatment since it is effective in detoxifying the free radicals induced by laser exposure. Our data suggests that the unexplored hypoxia/ROS/glutathione signaling axis may act as an important factor in the progressive pathological development and laser-resistant phenotypes of PWB, which will be the focus of our next studies.

There are several limitations to this study. First, metabolomic profiles from PWS lesions are yet to be determined. It is unknown whether glutathione and sphingolipid metabolism pathways are perturbed in PWB lesions. The upregulated HIF-1α and several key factors regulating glutathione metabolism indicate an elevation of glutathione in PWB lesions. However, the expression patterns of enzymes responsible for sphingosine metabolism are yet to be examined in PWB lesions. Second, sample size was limited; a follow-up study with a larger sample size including both biological and experimental replications should be conducted. Third, non-targeted metabolomics covers a wide range of metabolites, but absolute quantification of metabolites is lacking. A follow-up targeting metabolomic study for glutathione and sphingolipid metabolism is needed to validate the current study. Fourth, the exact link between perturbed metabolites and pathology in lesions is yet to be determined, which will be our next focus.

In summary, we have found that sphingosine and glutathione are two main perturbed metabolites in PWB iPSCs. PWB vasculature is under persistent and mild hypoxia. Nevertheless, our current data demonstrate that there are dysregulated cellular redox homeostasis and sphingolipids-mediated signalosomes in the PWB vasculature, which may facilitate cell survival and pathological progression. Furthermore, the elevated glutathione levels may contribute to laser-resistant phenotypes in the PWB vasculature.

