## supplementary files for "Perturbations of glutathione and sphingosine metabolites in Port Wine Birthmark patient-derived induced pluripotent stem cells"

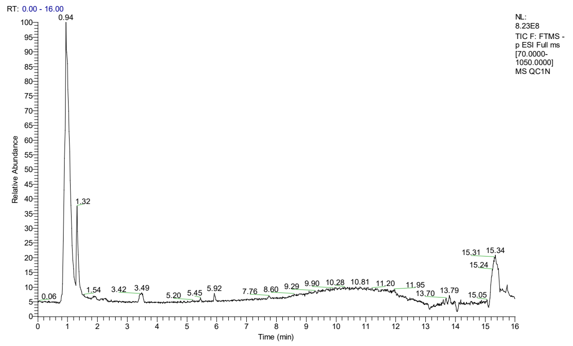


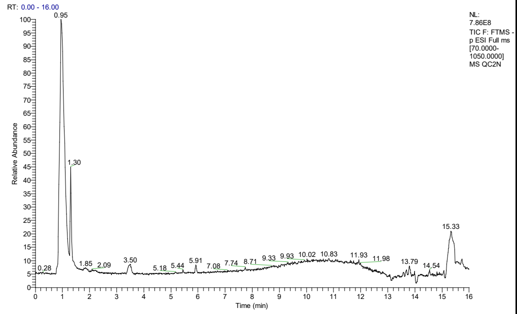


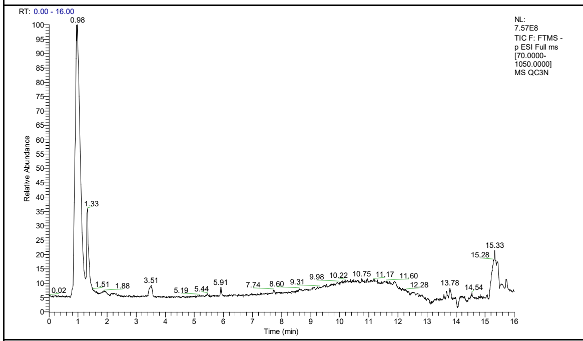


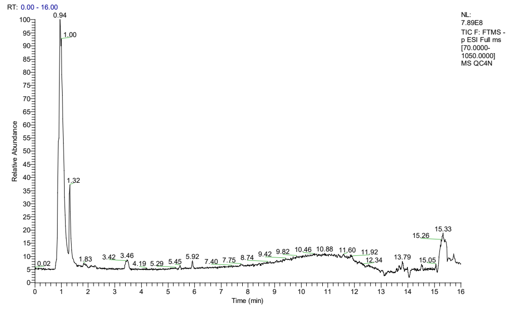


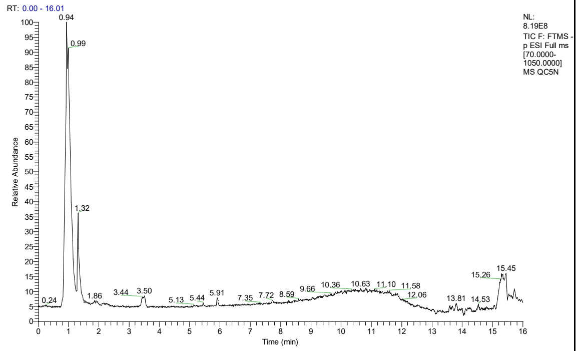


**Suppl. Fig 1A, TIC of metabolome analysis by LC-MS in ESI – mode; QC, n=5**

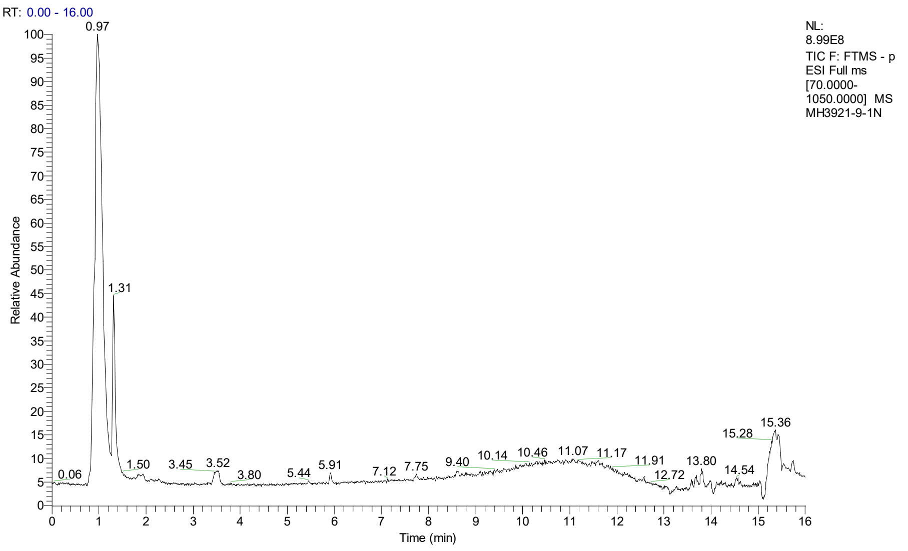


**Suppl. Fig 1 B, TIC for PWB_3921_9 in ESI - mode.**

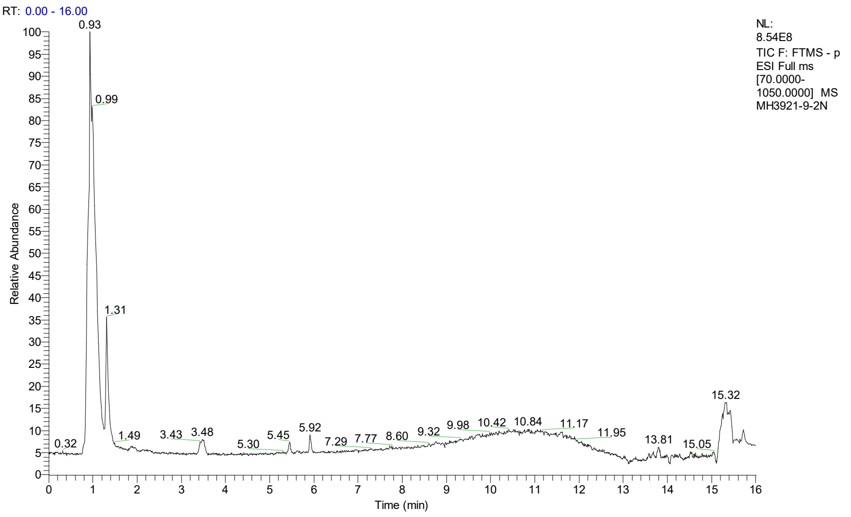


**Suppl. Fig 1 C, TIC for PWB_3921_9d in ESI - mode.**

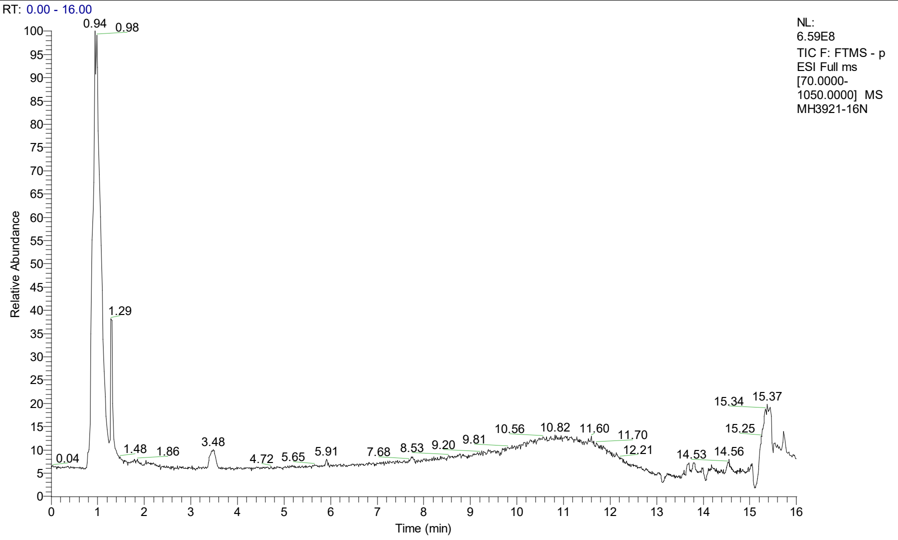


**Suppl. Fig 1 D, TIC for PWB_3921_16 in ESI - mode.**

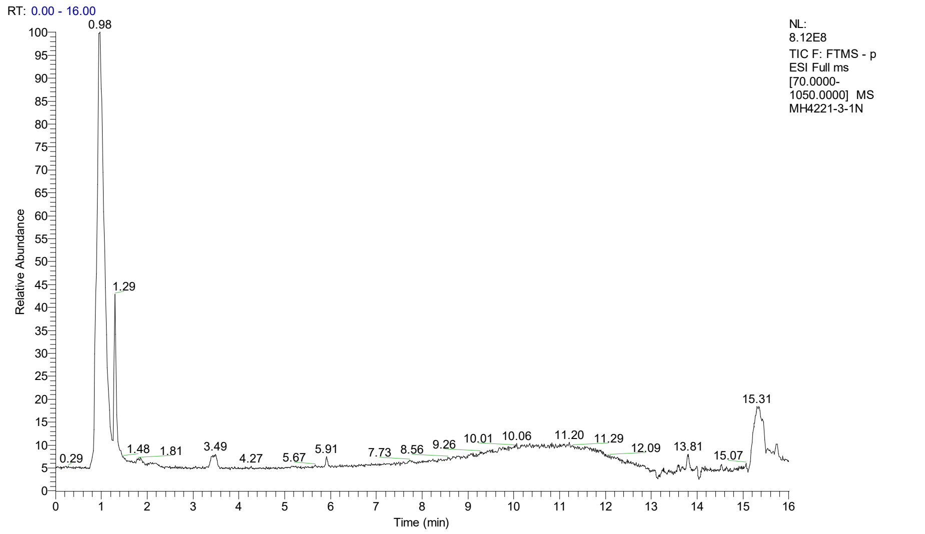


**Suppl. Fig 1 E, TIC for PWB_4221_3 in ESI - mode.**

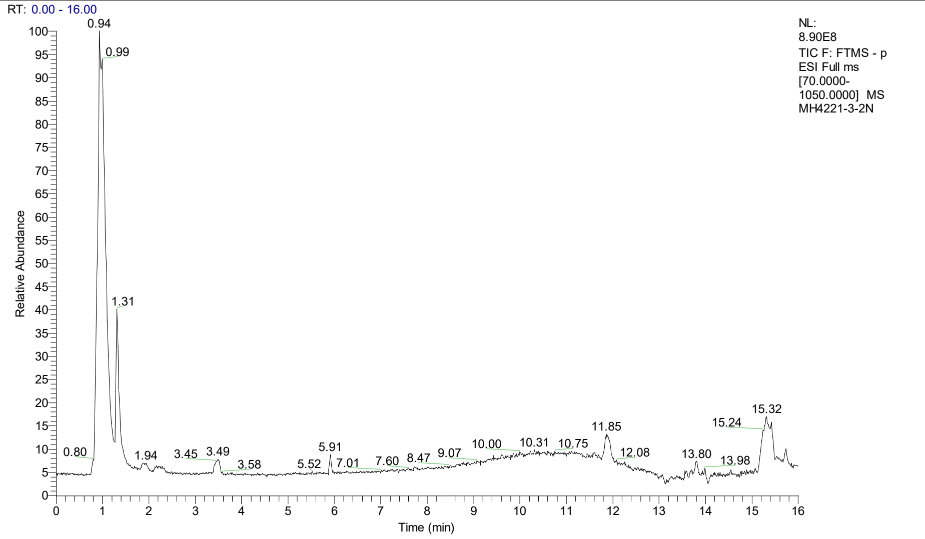


**Suppl. Fig 1 F, TIC for PWB_4221_3d in ESI - mode.**

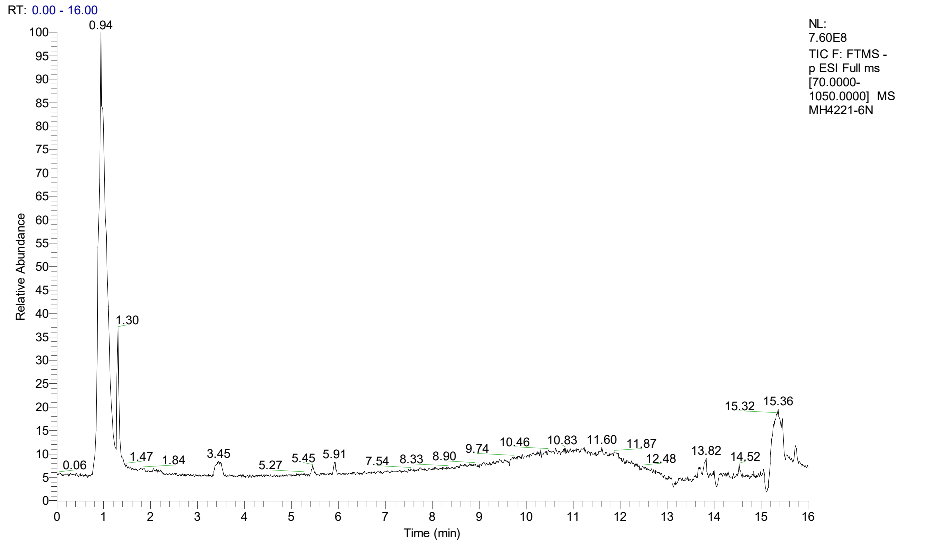


**Suppl. Fig 1 G, TIC for PWB_4221_6 in ESI - mode.**

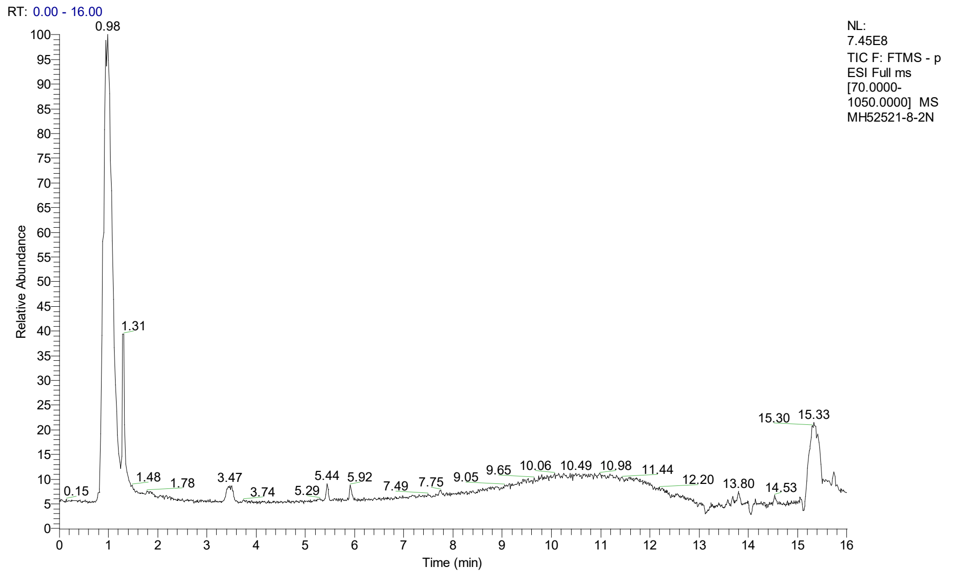


**Suppl. Fig 1 H, TIC for PWB_52521_8 in ESI - mode.**

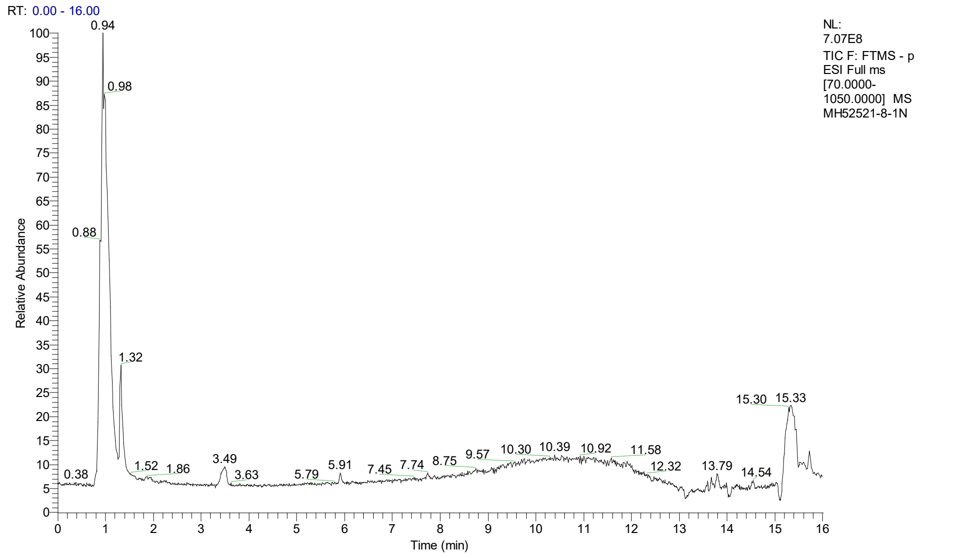


**Suppl. Fig 1 I, TIC for PWB_52521_8d in ESI - mode.**

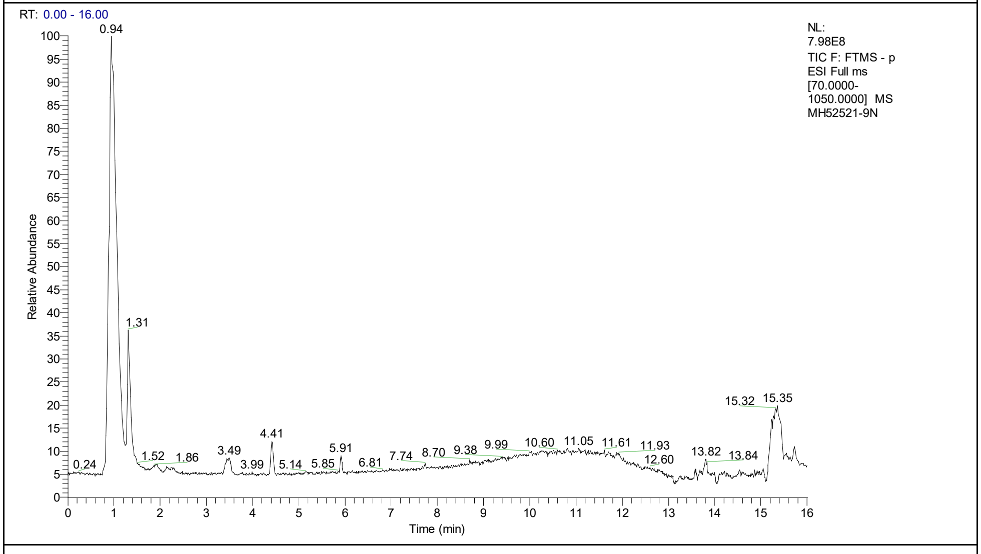


**Suppl. Fig 1 J, TIC for PWB_52521_9 in ESI - mode.**

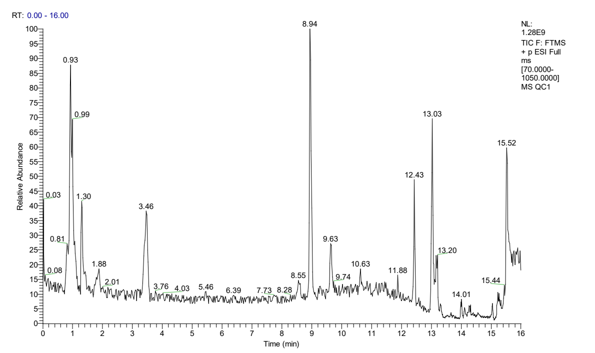


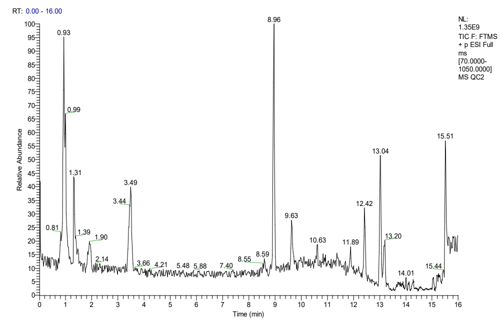


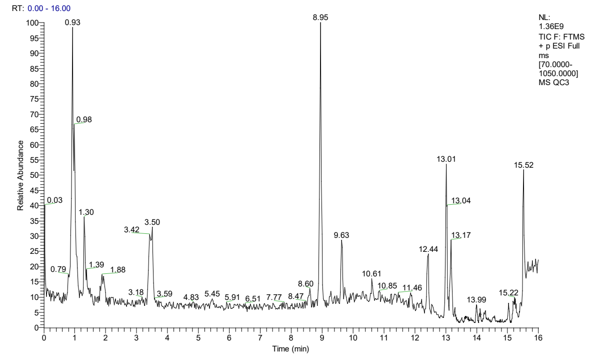


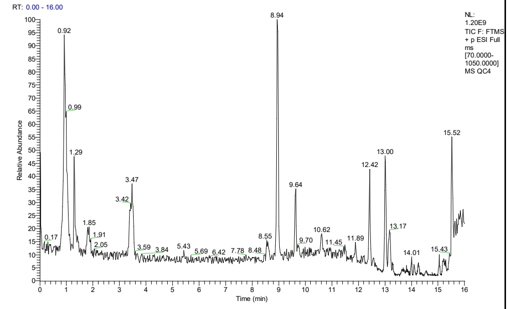


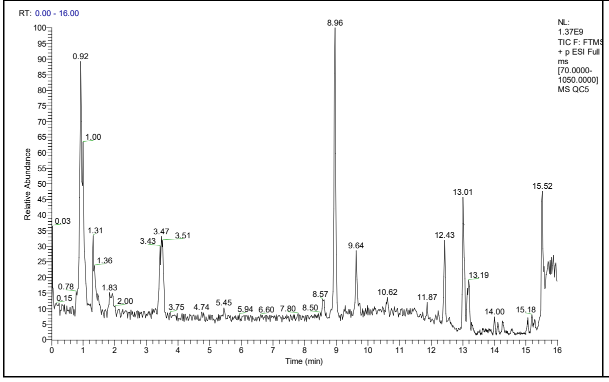


**Suppl. Fig 2A, TIC of metabolome analysis by LC-MS in ESI + mode; QC, n=5**

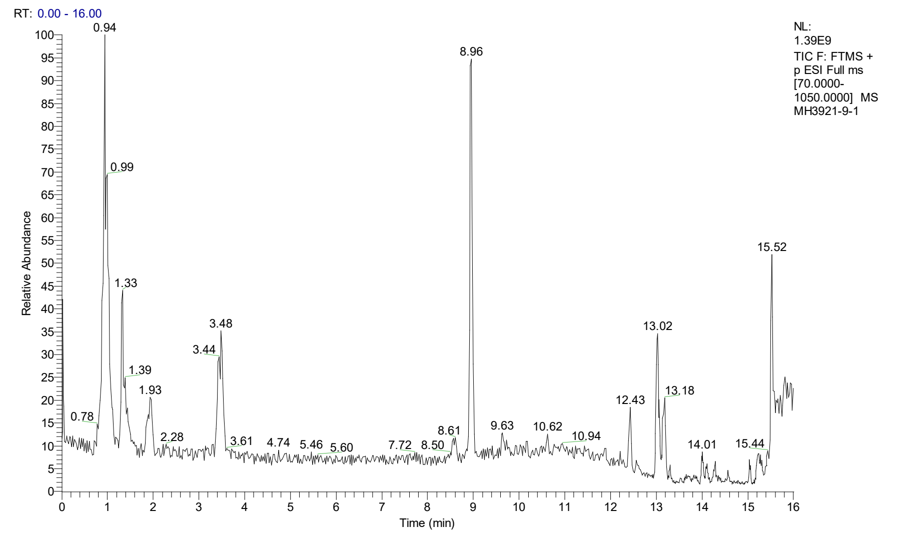


Suppl. Fig 2 B, TIC for PWB_3921_9 in ESI + mode.

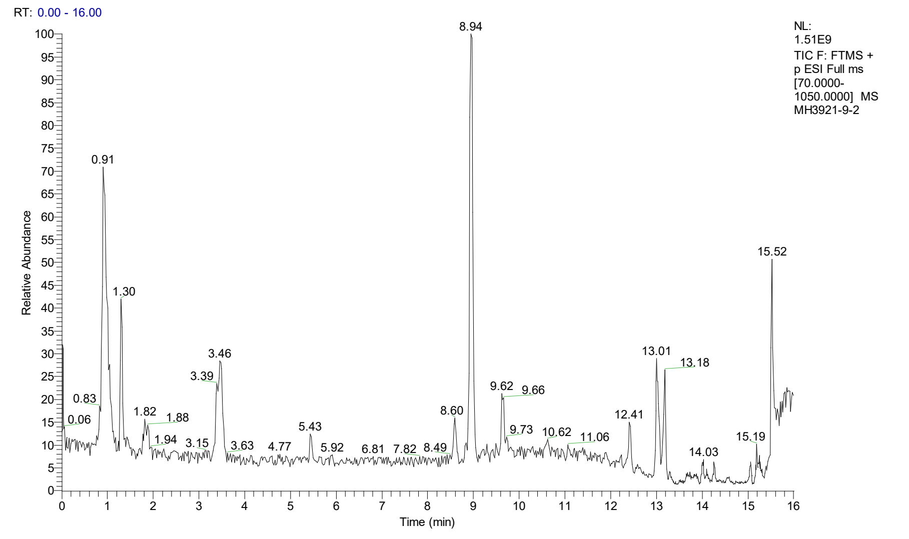


Suppl. Fig 2 C, TIC for PWB_3921_9d in ESI + mode.

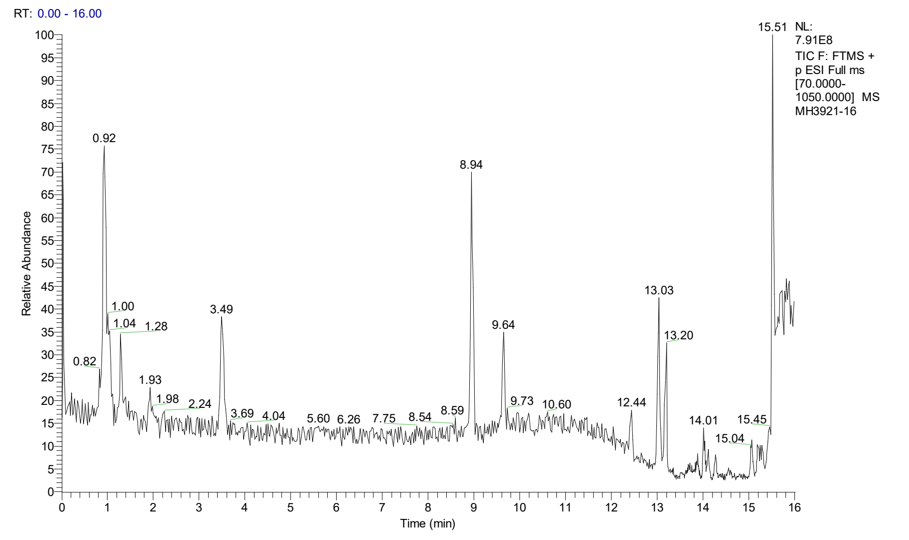


Suppl. Fig 2 D, TIC for PWB_3921_16 in ESI + mode.

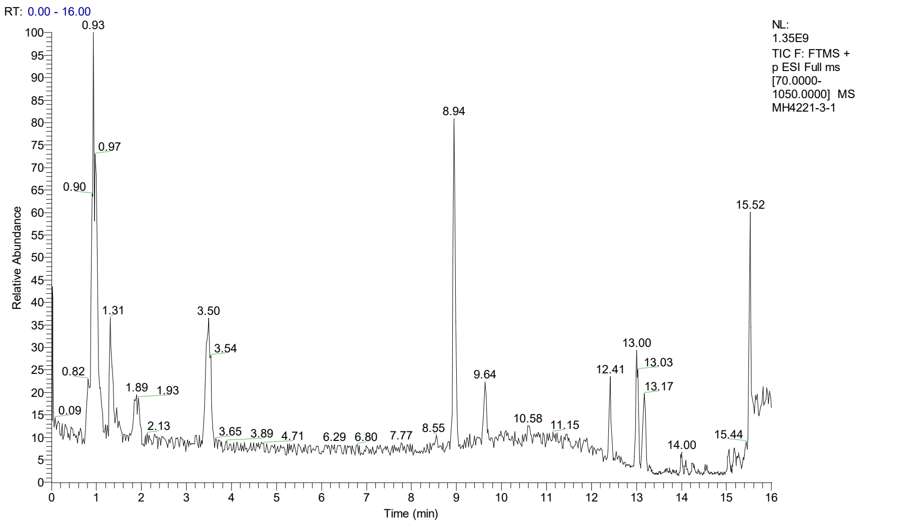


#### Suppl. Fig 2 E, TIC for PWB_4221_3 in ESI + mode.

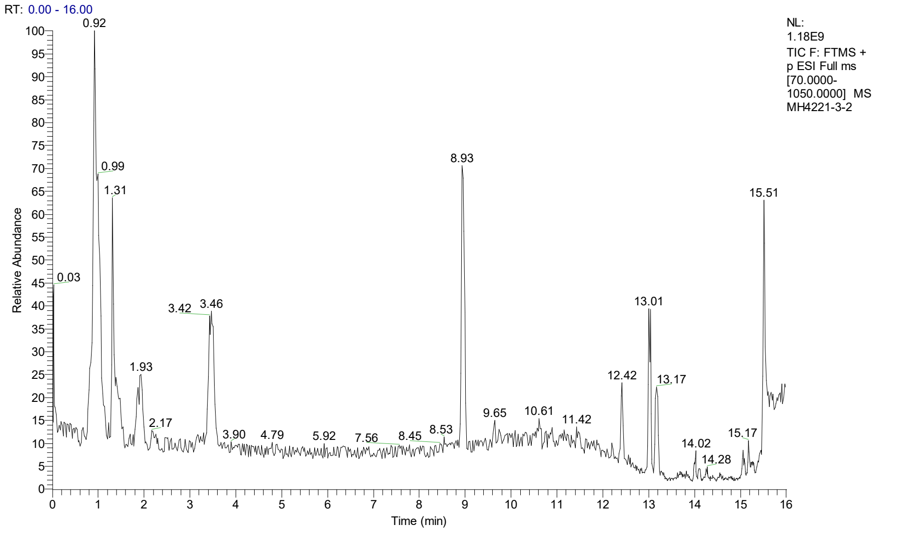


#### Suppl. Fig 2 F, TIC for PWB_4221_3d in ESI + mode.

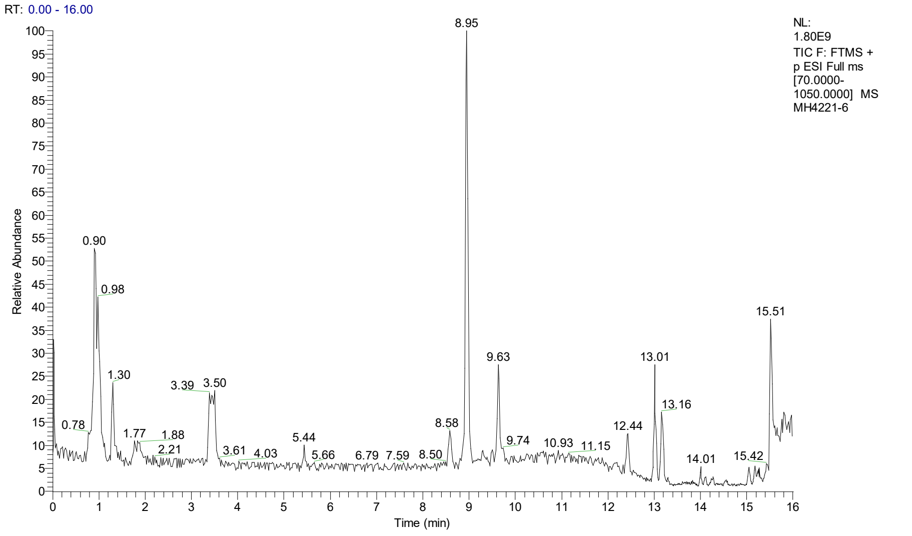


#### Suppl. Fig 2 G, TIC for PWB_4221_6 in ESI + mode.

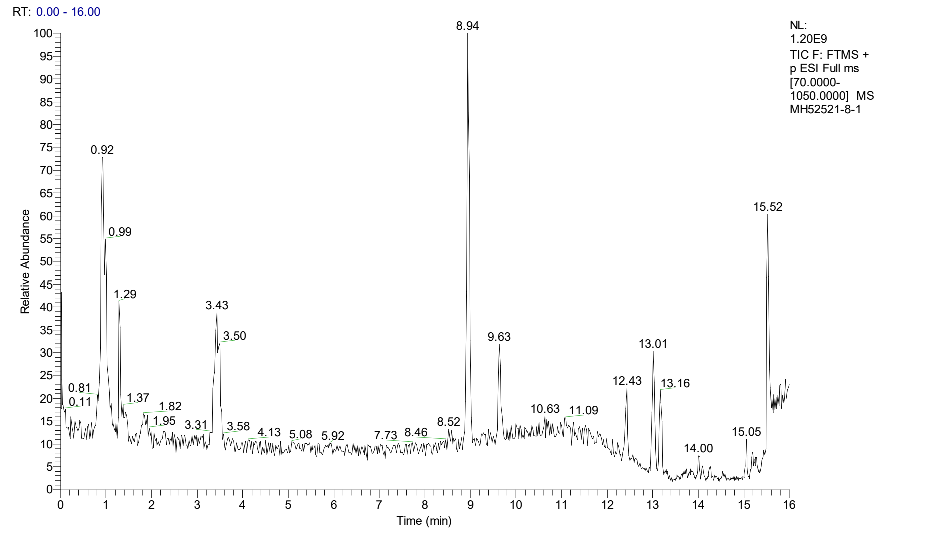


Suppl. Fig 2 H, TIC for PWB_52821_8 in ESI + mode.

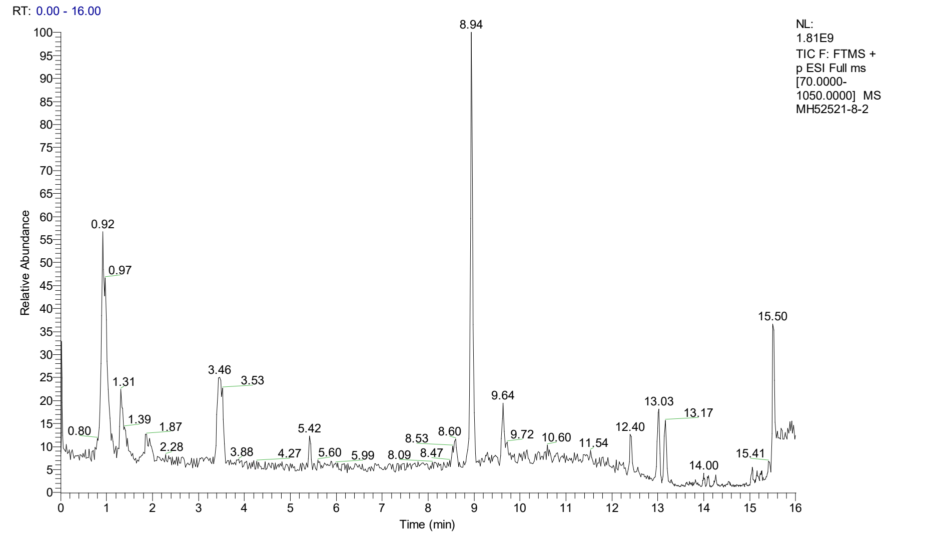


Suppl. Fig 2 I, TIC for PWB_52821_8d in ESI + mode.

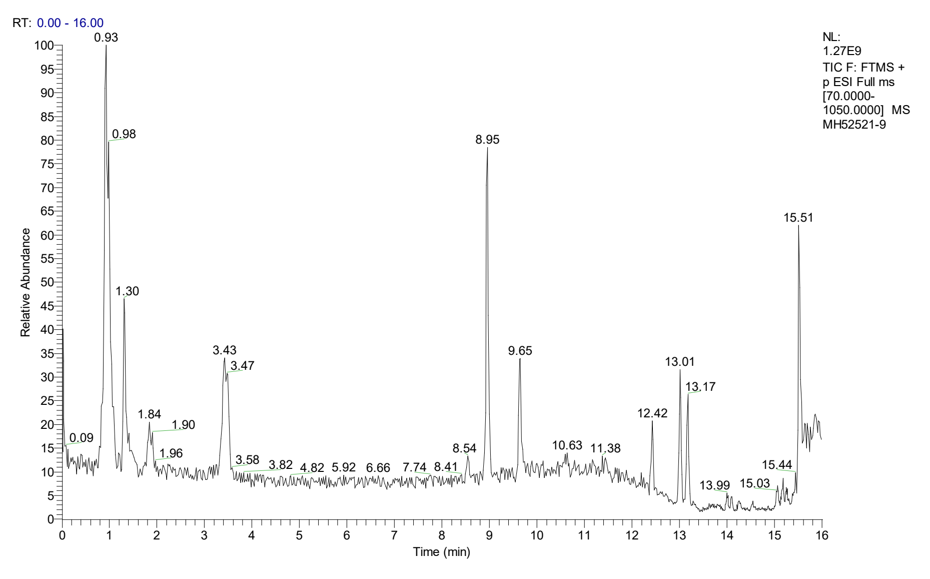


Suppl. Fig 2 J, TIC for PWB_52821_9 in ESI + mode.

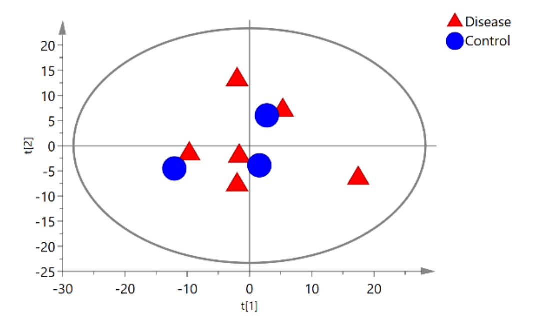


**Suppl Fig 3A. The scores scatter plot of PCA model in ESI – mode.**

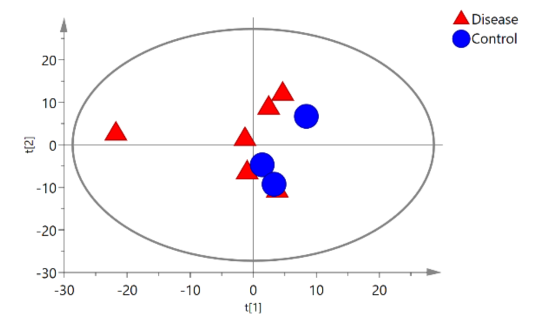


**Suppl Fig 3B. The scores scatter plot of PCA model in ESI + mode.**

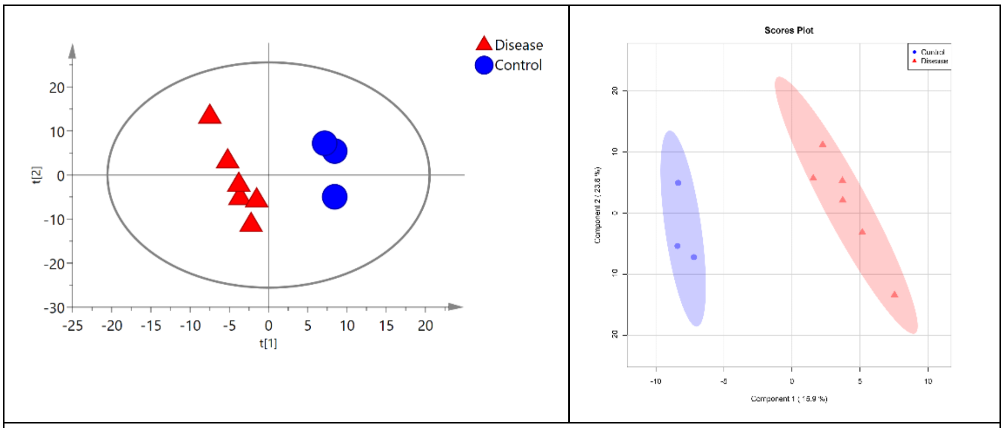


**Suppl Fig 4A. The scores scatter plot of PLS-DA model in ESI – mode.**

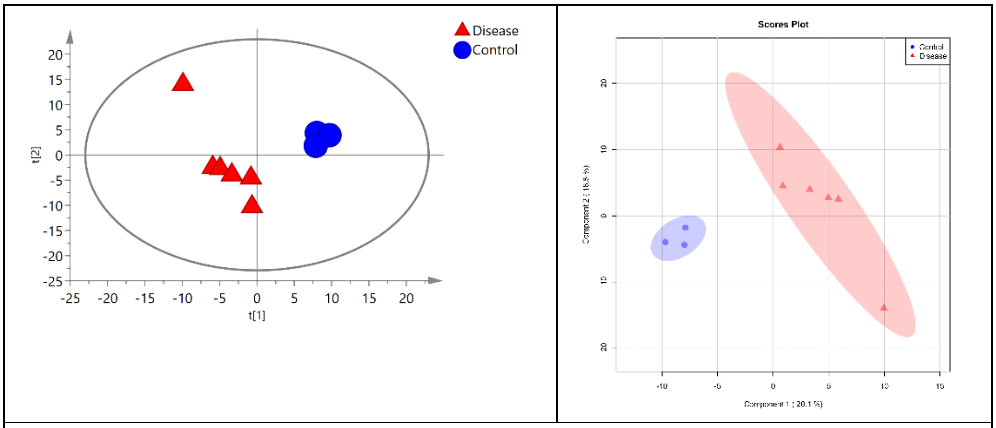


**Suppl Fig 4B. The scores scatter plot of PLS-DA model in ESI + mode.**

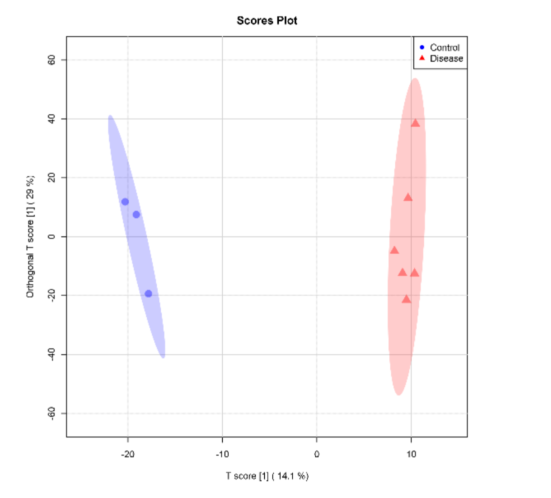


**Suppl Fig 5A. The scores scatter plot of OPLS-DA model in ESI – mode.**

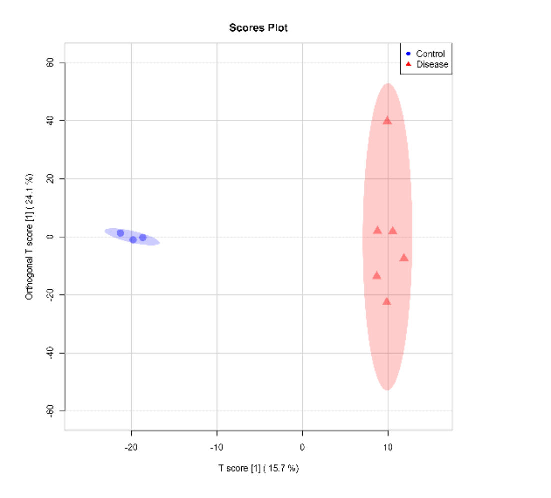


**Suppl Fig 5B. The scores scatter plot of OPLS-DA model in ESI + mode.**

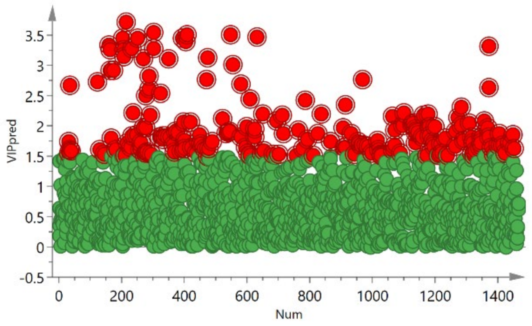


**Suppl Fig 6A. The loading plots of PLS-DA model in ESI – mode. Red box: metabolites with VIP > 1.5.**

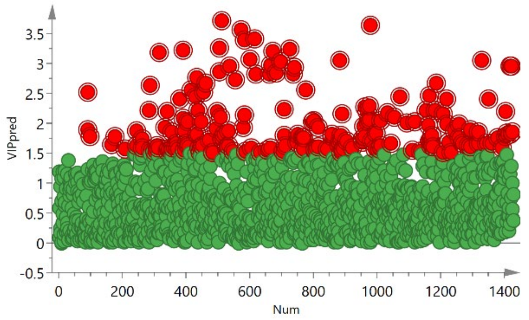


**Suppl Fig 6B. The loading plots of PLS-DA model in ESI + mode. Red box: metabolites with VIP > 1.5.**

#### Supplementary Table 1: The profile of the fragments identified in Fig 3

| Metabolites | Fragment formula | Fragment m/z | Peak m/z | Error (ppm) | Charge | Neutral charge |
|---|---|---|---|---|---|---|
| Sphingolipid_ESI+ | C4H9-e | 57.0699 | 57.0699 | 0.99 | 1 | |
| | C2H6NO-e | 60.0444 | 60.0445 | 1.81 | 1 | |
| | C5H11-e | 71.0856 | 71.0850 | -7.34 | 1 | |
| | C4H7NO+H | 86.0601 | 86.0598 | -3.49 | 1 | |
| | C6H10O-e | 97.0648 | 97.0645 | -2.37 | 1 | -H |
| | C6H10NO+H | 114.0914 | 114.0914 | -0.13 | 1 | +H |
| | C17H32N+H | 252.2686 | 252.2702 | 6.45 | 1 | +H |
| | C18H34N-e | 264.2686 | 264.2683 | -1.03 | 1 | |
| | C17H34NO+H | 270.2792 | 270.2769 | -8.25 | 1 | +H |
| | C18H36NO-e | 282.2792 | 282.2784 | -2.51 | 1 | |
| | C18H37NO2+H | 300.2897 | 300.2887 | -3.37 | 1 | |
| Glutathione_ESI+ | C2H4NO2+H | 76.0393 | 76.0389 | -5.42 | 1 | +H |
| | C4H6NO-e | 84.0444 | 84.0442 | -1.75 | 1 | |
| | C3H2NO2+H | 85.0159 | 85.0159 | 0.72 | 1 | |
| | C5H8NO3-e | 130.0499 | 130.0497 | -0.83 | 1 | |
| | C5H6NO2S-e | 144.0114 | 144.0114 | 0.11 | 1 | |
| | C5H8NO3S-e | 162.0220 | 162.0216 | -1.94 | 1 | |
| | C5H9N2O3S+H | 179.0485 | 179.0477 | -4.09 | 1 | +H |
| | C8H11N2O3S-e | 215.0485 | 215.0487 | 1.18 | 1 | |
| | C8H13N2O4S-e | 233.0591 | 233.0577 | -5.89 | 1 | |
| | C9H13N2O4S-e | 245.0591 | 245.0582 | -3.67 | 1 | |
| | C10H15N2O6S-e | 291.0646 | 291.0646 | 0.92 | 1 | |
| | C10H17N3O6S+H | 308.0911 | 308.0895 | -5.09 | 1 | |
| Glutathione_ESI- | C2H4NO2-H | 72.0091 | 72.0090 | -1.47 | 1 | |
| | C2H4NO2+e | 74.0247 | 74.0249 | 2.70 | 1 | |
| | C3H6NO2-H | 86.0247 | 86.0248 | 0.84 | 1 | -H |
| | C5H6NO2+e | 112.0404 | 112.0406 | 1.32 | 1 | |
| | C5H6NO3+e | 128.0353 | 128.0354 | 0.64 | 1 | |
| | C5H9N2O3-H | 143.0462 | 143.0464 | 0.93 | 1 | -H |
| | C5H8NO3S-H | 160.0074 | 160.0074 | 0.14 | 1 | -H |
| | C5H9N2O3S+e | 177.0339 | 177.0337 | -1.63 | 1 | |
| | C7H10N2O4-H | 185.0568 | 185.0565 | -1.57 | 1 | |
| | C9H12N3O5+e | 242.0782 | 242.0788 | 2.20 | 1 | |
| | C10H14N3O5-H | 254.0782 | 254.0777 | -2.19 | 1 | -H |
| | C10H16N3O6-H | 272.0888 | 272.0884 | -1.63 | 1 | -H |
| | C10H15N3O5S-H | 288.0659 | 288.0656 | -0.97 | 1 | |
| | C10H17N3O6S-H | 306.0765 | 306.0764 | -0.55 | 1 | |

**Suppl Fig 7 Original images for IHC.** Dashed blue boxed areas were used in the Fig 5 in each panel.

#### A. HIF-1α

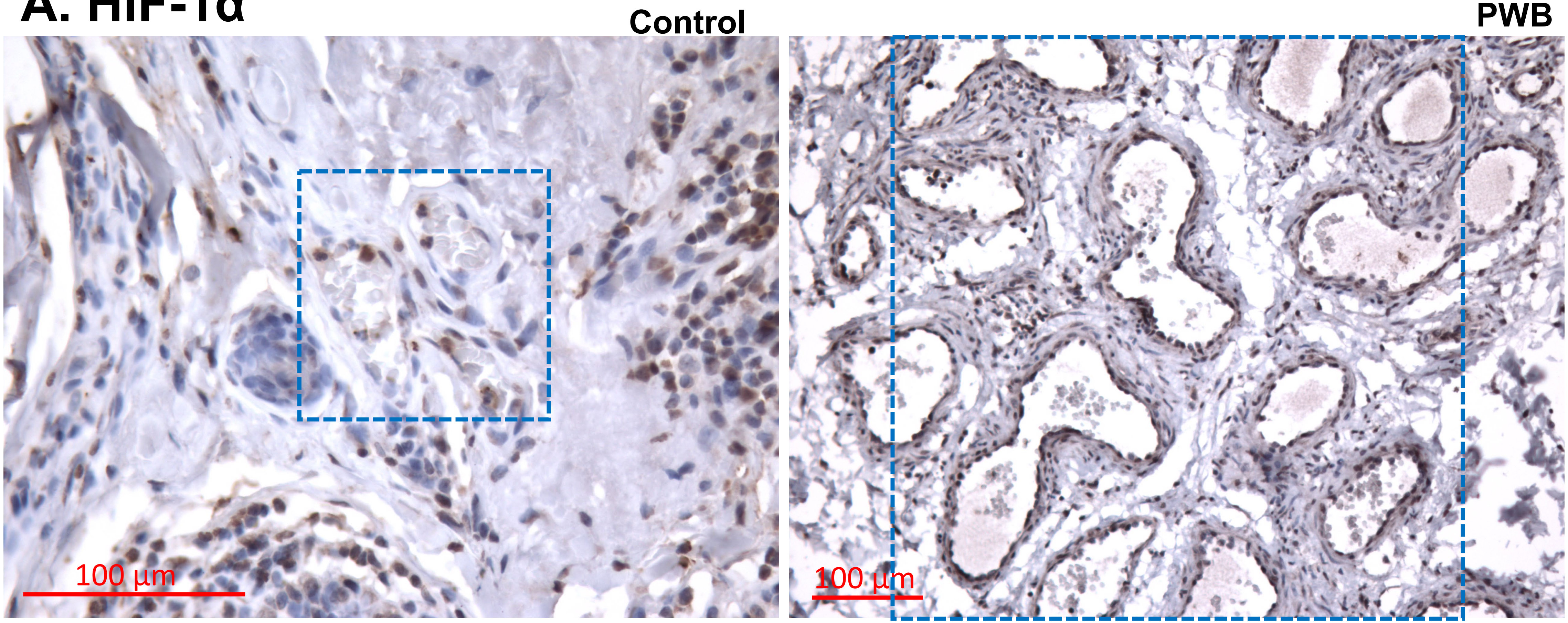

B. GCLM

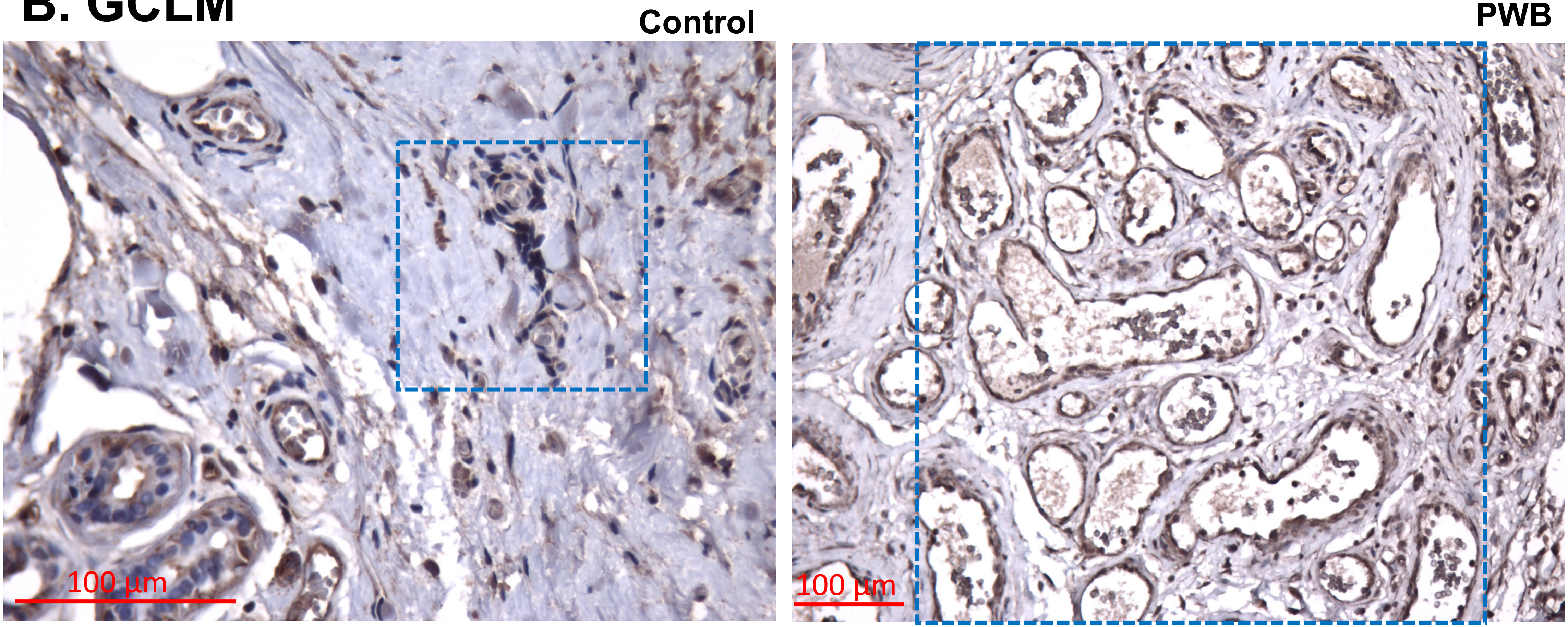

### C. GGT7

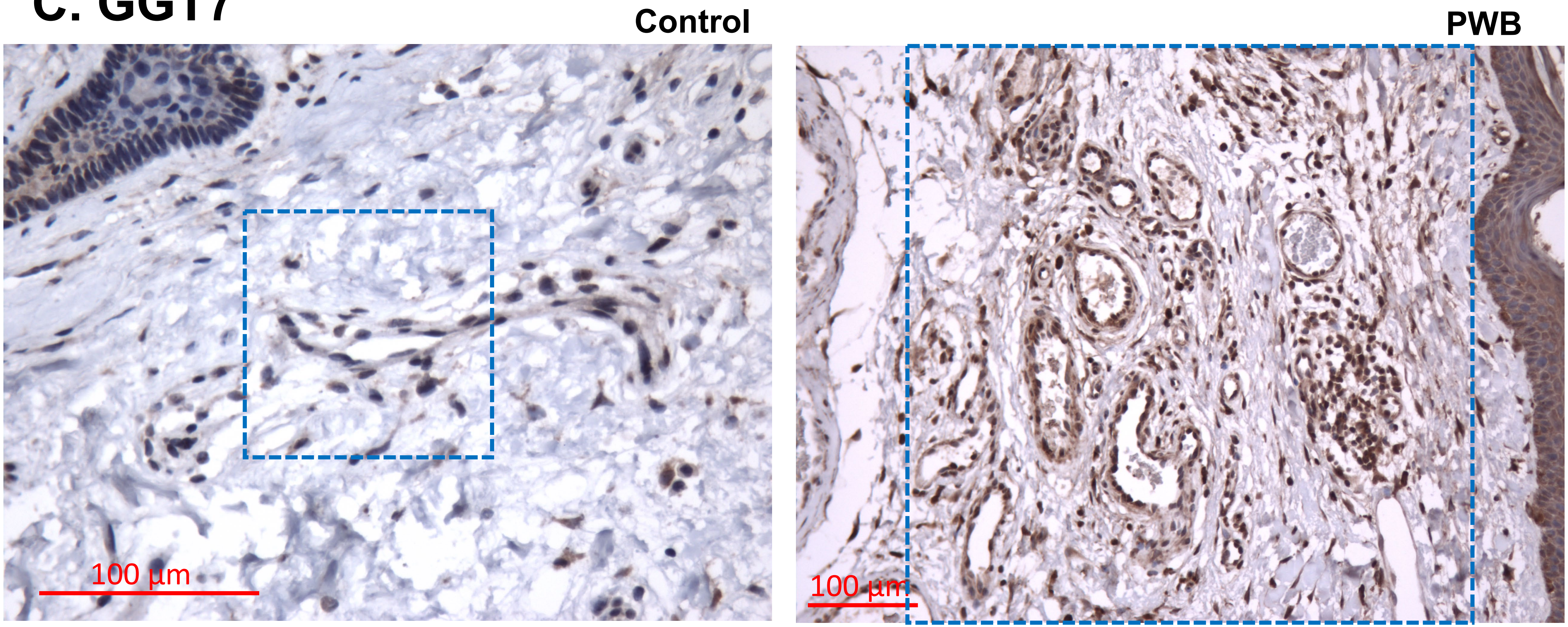

#### D. GSTP1

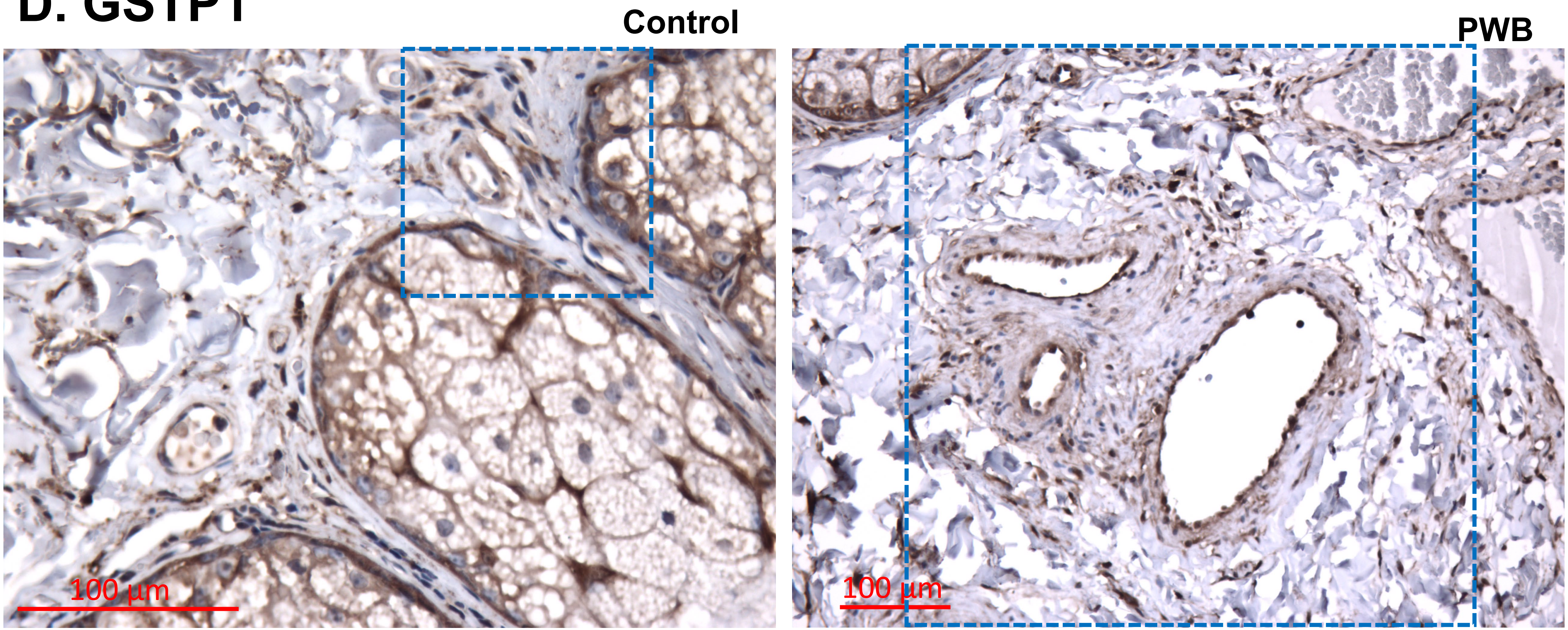
